# Immediate stimulus repetition abolishes stimulus expectation and surprise effects in fast periodic visual oddball designs

**DOI:** 10.1101/316786

**Authors:** Daniel Feuerriegel, Hannah A. D. Keage, Bruno Rossion, Genevieve L. Quek

## Abstract

Oddball designs are widely used to investigate the sensitivity of the visual system to statistical regularities in sensory environments. However, the underlying mechanisms that give rise to visual mismatch responses remain unknown. Much research has focused on identifying separable, additive effects of stimulus repetition and stimulus appearance probability (expectation/surprise) but findings from non-oddball designs indicate that these effects also interact. We adapted the fast periodic visual stimulation (FPVS) unfamiliar face identity oddball design (Liu-Shuang et al., 2014) to test for both additive and interactive effects of stimulus repetition and stimulus expectation. In two experiments, a given face identity was presented at a 6 Hz periodic rate; a different identity face (the oddball) appeared as every 7^th^ image in the sequence (i.e., at 0.857 Hz). Electroencephalographic (EEG) activity was recorded during these stimulation sequences. In Experiment 1, we tested for surprise responses evoked by unexpected face image repetitions by replacing 10% of the commonly-presented oddball faces with exact repetitions of the base rate face identity image. In Experiment 2, immediately repeated or unrepeated face identity oddballs were presented in high and low presentation probability contexts (i.e., expected or surprising), allowing assessment of expectation effects on responses to both repeated and unrepeated stimuli. Across both experiments objective (i.e., frequency-locked) visual mismatch responses driven by stimulus expectation were only found for oddball faces of a different identity to base rate faces (i.e., unrepeated identity oddballs). Our results show that immediate stimulus repetition (i.e., repetition suppression) can reduce or abolish expectation effects as indexed by EEG responses in visual oddball designs.

**Highlights:**

- We studied visual mismatch responses with a fast periodic oddball design
- Our design cleanly separates immediate stimulus repetition and expectation effects
- Stimulus expectation effects were only present for unrepeated stimuli
- Immediate stimulus repetition reduced EEG expectation effects

## 1. Introduction

The visual oddball design has been widely used to investigate the sensitivity of sensory systems to environmental statistical regularities (Grimm et al., 2016; Stefanics et al., 2014). In this design, a task-irrelevant critical stimulus is presented in high and low probability contexts. In the high probability context, a critical stimulus A is presented frequently as the ‘standard’ stimulus, interspersed with a rare ‘deviant’ stimulus B (e.g., AAAABAAABAAAAAAB… see Figure 1A). In the low probability context, the presentation probabilities for these stimuli are reversed, so that the original standard stimulus A is instead presented as a rare deviant (e.g., BBBBABBBABBBBBA… see Figure 1C). Comparisons of event-related potentials (ERPs) evoked in the human electroencephalogram (EEG) by the critical stimulus in the two contexts (i.e., A_Standard_ vs. A_Deviant_) reveal more negative-going waveforms evoked by deviants at posterior electrodes between 150-300ms (Czigler et al., 2004; Kimura et al., 2009; Stefanics, et al., 2011), an effect known as the visual mismatch negativity (vMMN; for recent reviews see Kimura et al., 2011; Stefanics et al., 2014). The vMMN is considered to be the visual counterpart of the earlier-discovered auditory MMN (Naatanen, et al., 1978), and has been similarly used to investigate a range of phenomena including sensory memory/change detection (Czigler et al., 2002), perceptual discrimination (Tales & Butler, 2006), stimulus repetition effects (Amado & Kovacs, 2016), and perceptual expectations (Stefanics et al., 2014). The magnitude of the visual mismatch response differs between healthy and clinical samples across a wide range of psychiatric and neurological disorders (reviewed in Kremlacek et al., 2016), as has also been found for the auditory MMN (reviewed in Naatanen et al., 2014).

**Figure 1.**
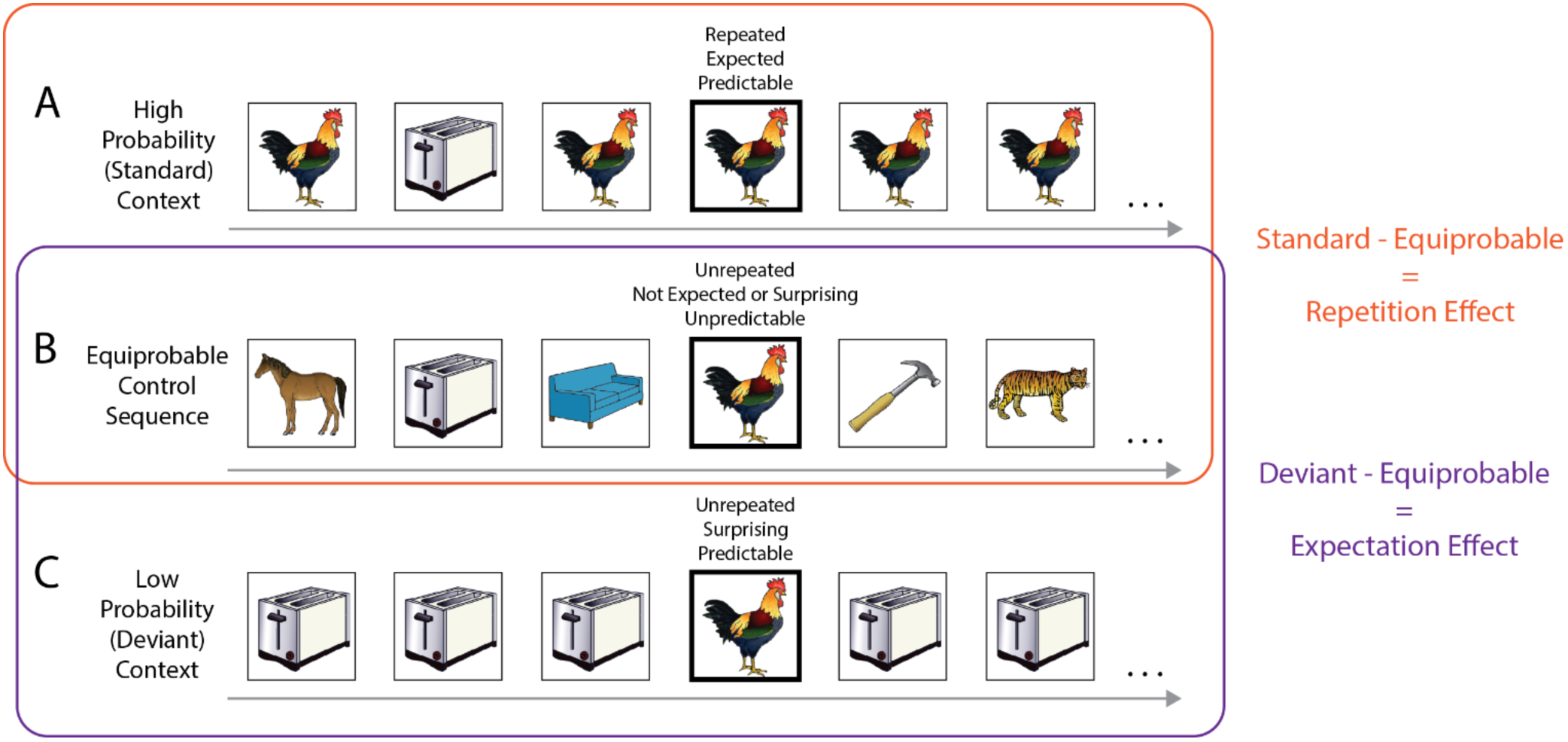
Examples of standard, deviant and equiprobable control sequences in a classical oddball design. The critical stimulus in each sequence is denoted by a thick black outline. A) In high probability (standard) sequences, a critical stimulus (e.g., rooster) is presented frequently, interspersed with a different rare stimulus (e.g., toaster). B) In equiprobable control sequences, the critical stimulus is presented rarely, within a sequence containing many different stimuli. C) In low probability (deviant) sequences, the critical stimulus appears with the same low probability as in equiprobable sequences, but is interspersed with another frequently-presented stimulus. Labels above each critical stimulus indicate whether it is *i)* immediately repeated/unrepeated, *ii)* expected/surprising, and *iii)* relatively predictable/unpredictable for the observer. Orange and purple rectangles and labels denote the sequences and comparisons used to derive stimulus repetition and expectation effects in previous studies. Example stimuli are used with permission from Rossion and Pourtois (2004).

The vMMN, or more generally the visual mismatch response (VMR), is thought to index effects of both *stimulus repetition* and *stimulus expectation* (often operationalised as the probability of a certain stimulus appearing). Stimulus repetition effects are commonly known as repetition suppression or stimulus-specific adaptation (Desimone, 1996; Movshon & Lennie, 1979) and are defined as a stimulus-specific reduction in a measure of neuronal activity (e.g., firing rate, local field potential amplitude, or fMRI BOLD signal change) in response to repeated compared to unrepeated stimuli (reviewed in Grill-Spector et al., 2006). Recent circuit models of repetition suppression (Dhruv et al., 2011; Solomon & Kohn, 2014; Kaliukhovich & Vogels, 2016) can explain standard/deviant ERP differences as reductions in stimulus-evoked responses to standards, due to mechanisms such as firing rate-dependent fatigue, synaptic depression or prolonged afterhyperpolarisation (Zucker & Regehr, 2002; Fioravante & Regehr, 2011, Vogels, 2016). This is accompanied by absent or reduced repetition suppression for deviants (May & Tiitinen, 2010; Nelken & Ulanovsky, 2007) or enhanced responses to deviants by adaptation-induced disinhibition of neurons responsive to deviant stimulus features (Solomon & Kohn, 2014; Kaliukhovich & Vogels, 2016).

On the other hand, VMRs have also been interpreted as expectation-driven effects. Theories of perception based on hierarchically-organised predictive coding (Friston, 2005; Rao & Ballard, 1999) posit that differences in standard and deviant responses result from larger prediction error signals to the rare and unexpected deviant stimuli (Friston, 2005; Garrido et al., 2009; Malmierca et al., 2015; Stefanics et al., 2014). Within the predictive coding framework, repetition suppression is conceptualised as a reduction of prediction error signals due to perceptual expectations that are weighted toward recently-encountered stimuli (Auksztulewicz & Friston, 2016; Summerfield et al., 2008). Stimulus repetition and expectation effects are both presumed to act on the same stimulus-selective neurons in the visual system, and may interact either each other as well as with other effects, such as attention, that modulate the same excitatory-inhibitory circuits (e.g., Reynolds & Heeger, 2009).

Importantly, the classical visual oddball design provides no way to disentangle the contributions of stimulus repetition and stimulus expectation. In these designs, the expected stimulus (the standard) is almost always a repeated stimulus, where the unexpected stimulus (the deviant) is never a repeated stimulus (compare Figures 1A, 1C). Separating the influences of each effect is critical to understanding how each modulation of neural activity separately and interactively facilitates discrimination between recently-seen and novel stimuli (i.e., change detection) and enables tracking of statistical regularities in sensory environments.

Recent studies have attempted to overcome the limitations of the classical oddball design, controlling for stimulus repetition effects in order to isolate the contribution of stimulus expectation. Such designs include additional sequences in which many different stimulus images (including the same stimuli used as standards and deviants in classical oddball sequences) are interspersed randomly within a sequence (known as equiprobable control sequences, e.g., Jacobsen & Schroger, 2001; Figure 1B). In these sequences, the critical stimulus has the same probability of appearance as it does in the deviant context within the classical oddball design. ERPs to the same stimulus are compared across equiprobable and standard contexts to test for repetition effects, and across equiprobable and deviant contexts to test for expectation/surprise effects (e.g., Amado & Kovacs, 2016; see sequence comparison labels in Figure 1). Results of these studies already suggest that both stimulus repetition effects and expectation effects contribute to the VMR (Amado & Kovacs, 2016; Astikainen et al., 2008; Czigler et al., 2002; Kimura et al., 2009). However, several important confounds persist when using equiprobable control sequences, which can be understood by considering the types of expectations that can be formed during each sequence type. In classical oddball sequences (i.e., those in Figures 1A, 1C), expectations can be formed specifically for the repeating standard stimulus image, such that surprise responses can be elicited by deviant stimuli which violate such image-specific expectations. However, stimulus image-specific expectations cannot be formed (or violated) for the larger number of randomly-interspersed stimuli in the equiprobable sequences. Comparing the same stimulus across standard and equiprobable contexts (to measure repetition effects) therefore also involves comparing an expected stimulus (standard) with a stimulus that is neither expected nor surprising (equiprobable). Such a comparison does not in fact manage to isolate repetition effects, since it necessarily confounds stimulus expectation with stimulus repetition. Additionally, since standard/deviant sequences only ever contain a maximum of two stimuli, after a brief period of exposure these two specific stimulus images become readily predictable for participants. By contrast, the large number of possible stimuli that appear in equiprobable sequences (usually ~10) make the image properties of any individual stimulus comparatively unpredictable. Note that here we use the term *predictability* to refer to the *range* of possible stimuli that could appear, rather than the expectation that a specific stimulus image will appear next. Given that effects of stimulus repetition, expectation/surprise, and stimulus predictability have all been found on visual stimulus evoked ERPs within the time range of the VMR (Feuerriegel et al., 2018), it is unclear how each mechanism may have contributed to effects observed when using comparisons with equiprobable sequences. To tease these separate aspects apart, a design that independently manipulates stimulus repetition and expectation without a stimulus predictability-related confound is required, but is missing from the literature on the VMR thus far.

Not only are stimulus repetition and expectation effects readily confusable, evidence from other (non-oddball) experiments also indicates that these effects may actually interact. Recent work has found that repetition effects are larger for surprising stimuli, due to large surprise-related signal increases for unrepeated stimuli (Amado & Kovacs, 2016; Larsson & Smith, 2012; Choi et al., 2017; Kovacs et al., 2012, 2013; reviewed in Kovacs & Vogels, 2014). Similar interaction effects have also been found for ERPs/ERFs, with several studies showing that expectation violation responses can be reduced when stimuli are repeated (Todorovic & de Lange, 2012; Symonds et al., 2017; Wacongne et al., 2011). Demonstrating similar interactions in visual oddball designs is critical to extending existing models of novelty/change detection (e.g., Kremlacek et al., 2016) to incorporate interacting mechanisms driven by recent stimulus exposure (stimulus repetition) and longer-term stimulus appearance probabilities (expectation). Where such models have already been partially developed for the auditory mismatch response (see Costa-Faidella, et al., 2011; Mittag et al., 2016), similar interacting mechanisms have not yet been characterised in the visual domain.

The goal of the present study was to develop a visual oddball design that cleanly separates stimulus repetition and expectation effects while controlling for stimulus predictability, our primary aim being to test for additive and interactive effects of stimulus repetition and expectation. To this end, we developed a highly-efficient design that can present both repeated and unrepeated stimuli in expected and surprising contexts, can present many rare deviant stimuli in a short period of time to elicit high signal-to-noise ratio (SNR) responses, and allow objective identification of the presence of a response – and its quantification – in the EEG frequency domain. Our design is based on a recently-developed oddball design in the context of Fast Periodic Visual Stimulation (FPVS; Liu-Shuang et al., 2014, 2016; Dzhelyova & Rossion, 2014; Dzhelyova et al., 2017). In the FPVS oddball design, a base rate stimulus is presented at a rapid, periodic rate (e.g., 6 Hz). Oddball stimuli replace the base rate stimulus every *N* stimuli at a fixed periodicity (e.g., 1/7 stimuli = 0.857 Hz oddball periodicity). EEG frequency domain responses to these stimulation sequences exhibit high SNR responses that are robust against non-periodic artefacts such as blinks and motor responses (Liu-Shuang et al., 2014; Dzhelyova et al., 2017). Moreover, these responses can be identified objectively (i.e., at frequencies known in advance) and quantified in the frequency domain, avoiding more subjective evaluation of time-domain waveforms for the presence or absence of a given ERP component. Critically, responses at the frequencies of oddball stimulation reflect the extent to which the waveforms evoked by oddball and base rate stimuli differ, specifically indexing the *differential responses* between these stimulus types in the brain. Time-domain waveforms aligned to oddball stimulus onset can also be analysed to examine the time course of oddball-evoked responses (Dzhelyova & Rossion, 2014; Dzhelyova et al., 2017).

Here we adapted the FPVS oddball design to test for effects of stimulus expectation (independently of stimulus repetition) in Experiment 1, and for additive and interactive effects of immediate stimulus repetition and expectation in Experiment 2. In both experiments, we presented face images at a presentation rate of exactly 6 Hz, with an oddball face stimulus presented as every 7^th^ stimulus. Faces are an ideal stimulus type for our purposes here, since they are associated with robust face identity repetition and expectation effects (Henson, 2016; Grotheer and Kovacs, 2014;Summerfield et al., 2018; Feuerriegel et al., 2018). Moreover, in FPVS oddball designs, faces elicit a complex EEG response to, for example, changes of identity or facial expression (e.g., Dzhelyova & Rossion, 2014; Dzhelyova et al., 2017, respectively). In our FPVS oddball design observers can predict *when* an oddball will appear (e.g., after 6 base rate stimuli have appeared). By manipulating the presentation probability (i.e., perceptual expectations) for different oddball stimulus images within a sequence, we could entrain expectations relating to *which type* of oddball stimulus will appear. This allowed us to cleanly separate and quantify the influences of immediate stimulus repetition and stimulus expectation/surprise, which has not been possible using existing visual oddball designs.

## 2. Experiment 1

The goal of Experiment 1 was to test for surprise responses to unexpected stimuli in the visual oddball paradigm that could not be accounted for by stimulus repetition effects. In each stimulation sequence, we presented base rate faces at a presentation rate of 6 Hz, with a single oddball face replacing the base rate face every 7 stimuli at a fixed periodicity of 0.857 Hz (Figures 2A, 2C). We facilitated participants’ expectations for a *specific* oddball face image (of a different identity to the base rate faces) by presenting the same oddball face identity 90% of the time. Critically, this common (i.e., expected) oddball was replaced by an exact image repetition of the base rate faces for a small proportion (10%) of oddballs, resulting in a repeated, but also rare and surprising stimulus (see Figure 2A, 2B). Our design differs from previous FPVS face identity oddball designs, in that we presented only two different face images as oddballs within any given sequence (entraining expectations for specific oddball images), whereas previous experiments presented a large range of different face identities as oddball stimuli within a sequence (e.g., Liu-Shuang et al., 2014, 2016; Dzhelyova & Rossion, 2014).

**Figure 2.**
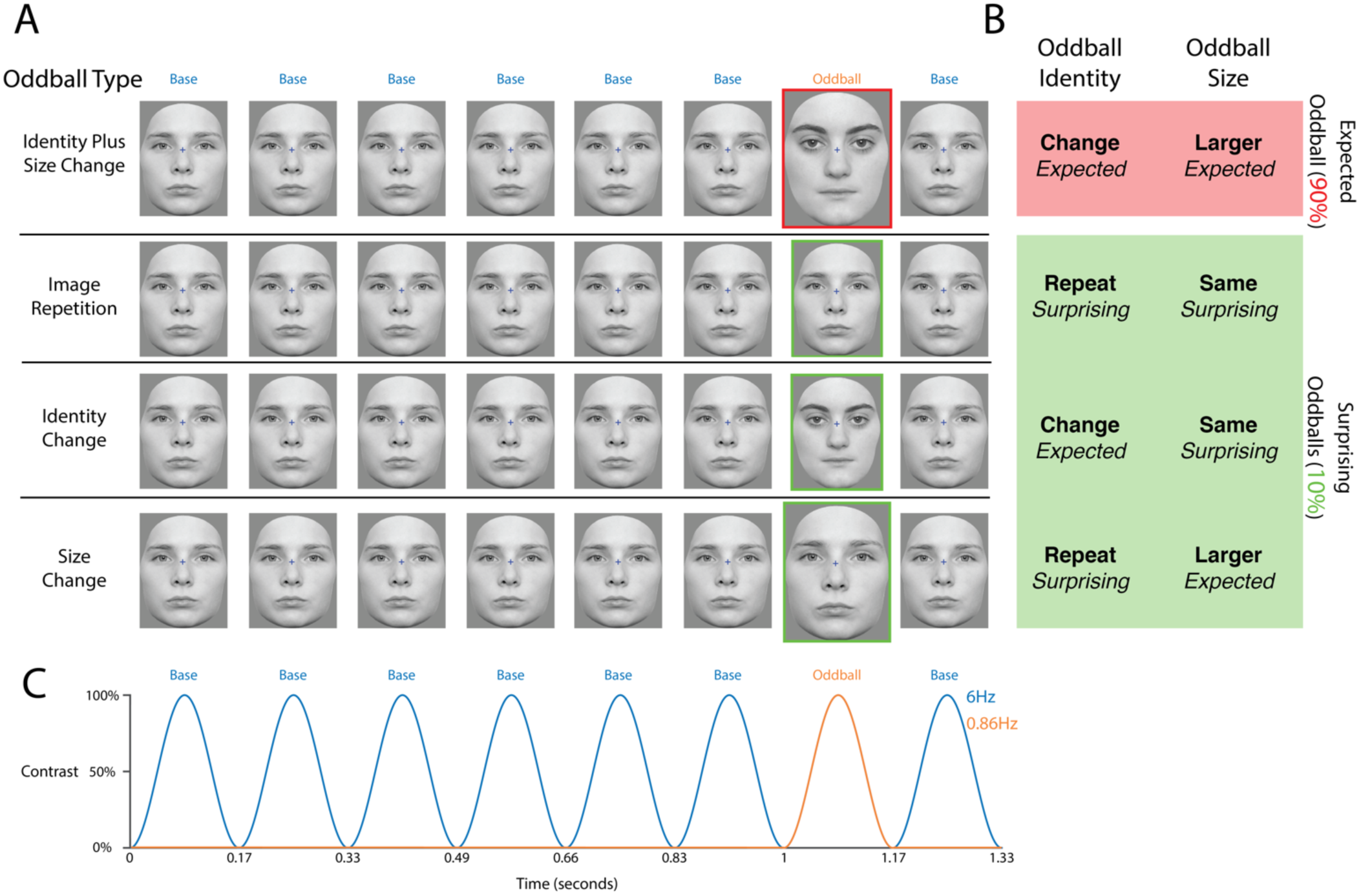
A) Explanation of oddball types in Experiment 1. Top Row: In each sequence a base face was presented at a rate of 6Hz. Within every sequence, 90% of oddballs were of a different identity to the base face and 20% larger in size - the so-called common oddball (the identity of this face did not change throughout the sequence). Thus participants could reliably expect every 7^th^ image to be both a different size and a different identity to the preceding base rate faces. Rows 2-4: The remaining 10% of oddballs were one of three “surprising” oddball types, which appeared in separate sequences. Each surprising oddball differed from the common oddball (and thus violated participants’ expectations) in either identity and/or size. B) Description of each oddball face’s identity and size relative to the base rate face (bold text) and the expected oddball (italicised text). C) Schematic of the sinusoidal contrast modulation used to achieve the 6 Hz presentation rate. Oddball faces (outlined in orange) appeared after every 6 base faces at a rate of 0.857 Hz.

We quantified EEG responses to these expected and surprising oddball instances by deriving 1-second epochs time-locked to oddball stimulus onset, and then concatenating these epochs to form continuous sequences (the so-called ‘false sequencing’ approach, see Quek & Rossion, 2017) with oddball responses occurring at a newly-imposed periodicity of 1-second (i.e., 1 Hz). For all oddball types, the periodic response at exactly 1 Hz (as measured in the frequency domain) reflects the degree to which the visual system distinguishes between base and oddball stimuli. For the surprising image repetition oddballs, a periodic response at the frequency of oddball stimulus presentation will selectively index stimulus expectation (anticipation of the common oddball stimulus appearance) that *cannot* be accounted for by stimulus repetition. Our FPVS design in this way allows us to test for the presence of expectation effects by assessing whether EEG responses at predefined oddball stimulation frequencies differ from zero. Based on previous reports of co-occurring repetition and expectation-related effects in visual oddball designs (e.g., Kimura et al., 2009), we expected to observe measurable oddball responses evoked by these surprising image repetition oddballs.

### 2.1 Participants

Twenty-two people participated in Experiment 1 (6 males, age range 18-27 years, mean age 21.7 ± 2.4 years). All participants had normal or corrected-to-normal vision and were right-handed as assessed by the Edinburgh Handedness Inventory (Oldfield, 1971). All participants showed unimpaired facial recognition ability as measured by the electronic version of the Benton Facial Recognition Test (Benton et al., 1983; see Rossion & Michel, 2018 for the electronic version; all global accuracy scores ≥ 39/54 cutoff defined in Rossion & Michel). This study was approved by the Biomedical Ethical Committee of the University of Louvain.

### 2.2 Stimuli

Thirty-six frontal images of faces (18 male, 18 female, neutral expression) were taken from the stimulus set in Laguesse and Rossion (2013). We converted images into greyscale and equated their mean pixel intensity and RMS contrast using the SHINE toolbox (Willenbockel et al., 2010). Stimuli subtended 8.4° × 8.6° visual angle at a viewing distance of 80cm. We also created a separate set of larger faces by scaling the resulting face images by 120%.

### 2.3 Procedure

Participants sat in a dimly-lit room 80cm in front of an LED monitor (refresh rate 120 Hz) and viewed stimuli presented against a grey background. A small fixation cross was superimposed over the nasion of the face images throughout the sequence. Each stimulation sequence consisted of a fixation cross for 2 seconds, followed by an 86-second stimulation period, and then another 2-second period during which only the fixation cross was visible. During the stimulation period, we used an in-house developed program (SinStim) to present faces at a rate of 6 Hz using a sinusoidal contrast modulation. In each 166.66ms image cycle, the stimulus image contrast was smoothly modulated from 0% to 100% to 0%. During the first 2 seconds of the stimulation period, the maximum contrast within a cycle gradually increased from 0-100% (i.e., a fade-in), and gradually decreased to 0% across the last 2 seconds of the stimulation period (i.e., a fade-out).

An overview of the stimulation sequences is given in Figure 2C. In each sequence a single identity base rate face appeared at a periodic rate of exactly 6 Hz. Within this sequence, an oddball face – of a different identity to the base rate faces - was presented as every 7^th^ image in the sequence (i.e., 6 Hz/7 = 0.857 Hz). This oddball face identity appeared as 90% of all oddballs within the sequence, thus we refer to it as the “common oddball”. The common oddball face was 20% larger than the base rate faces, and is accordingly labelled the <u>Identity Plus Size Change</u> oddball in Figure 2A. The face identity of this common oddball was the same face identity throughout a stimulation sequence. We anticipated that the reliable appearance of this common oddball every 7 images should lead participants to form expectations for this specific face identity/size combination, which, critically, we could then violate. Thus, for 10% of oddball presentations, we replaced the common (i.e., expected) oddball with a rare (i.e., surprising) oddball image, which differed across three sequence types.

Examples of each oddball type are displayed in Figure 2A. Here it is important to note that the oddball-specific EEG responses in our experiment index *differences* in responses to base rate and oddball stimuli, which can be due to both physical stimulus differences or expectation/surprise signals time-locked to oddball face presentation. For example, the signal evoked by the expected Identity Plus Size Change oddball could be driven by both low-level stimulus and face identity differences relative to the base rate faces (indexing differential visual responses and a release from repetition suppression), *and* also expectation signals time-locked to oddball onset. Stimulus differences between the base rate faces and each oddball type are listed in bold in Figure 2B. The expectation status of oddball stimulus attributes (with respect to the expected Identity Plus Size Change oddball) are listed in italicised text. Each oddball stimulus is named according to its stimulus characteristics with respect to the base rate faces.

In sequence type 1, the surprising oddball was the exact same image as the base rate face (i.e., an *unexpected repetition* of the base rate face, termed the <u>Image</u> <u>Repetition</u> oddball). As there are no physical stimulus differences from the base rate faces, any oddball-specific signals would be caused solely by expectation or surprise signals time-locked to oddball stimulus onset. We also included two other rare/surprising oddball types in separate sequences: *i)* the same face identity as the expected oddball, but the size of the base rate faces (termed the <u>Identity Change</u> oddball) in sequence type 2, and *ii)* an oddball of the same face identity as base rate faces, but the size of the expected oddball stimulus (termed the <u>Size Change</u> oddball) in sequence type 3. Since responses evoked by the Identity Change oddball index a combination of low-level and face identity differences relative to base rate faces (i.e., release from repetition suppression), we included this oddball type to estimate the magnitude of repetition suppression effects. In contrast, the Size Change oddball indexes a combination of low-level stimulus differences relative to base rate faces (due to increased image size) *and* surprise signals resulting from a violation of expectations for the common oddball face identity. This oddball (as well as the Identity Change oddball) were included to test the hypothesis that face identity repetition suppression (i.e., smaller signals for identity repetitions compared to identity changes from base rate faces) would not be found when identity repetitions are unlikely, indicating that repetition suppression effects simply reflect perceptual expectations (Summerfield et al., 2008; but see Pajani et al., 2017). We could test this hypothesis by comparing oddball responses between each size-matched identity repetition and identity change oddball stimulus type.

At least 5 expected oddballs were presented between each surprising oddball (mean = 9, range = 5-13). We presented 6 sequences per sequence type (total of 18 sequences), and 7 surprising oddballs in each sequence for a total of 42 surprising oddballs per sequence type. Within a single sequence, we presented faces of the same sex as base and oddball stimuli, resulting in sequences of only female or only male faces (3 male and 3 female sequences per oddball sequence type). Additionally, we counterbalanced the face identities allocated to each sequence type across participants. Total testing duration was 27 minutes.

### 2.4 Experimental Task

We used an orthogonal task to engage participants’ attention throughout the experiment (see Liu-Shuang et al., 2014; Rossion & Boremanse, 2011). Participants fixated on a blue fixation cross overlaying the images and pressed the spacebar when it changed colour from blue to red (10 colour changes per sequence, 100ms colour change duration, >2 seconds between colour changes). We considered key presses within 1000ms of a fixation cross colour change as correct.

### 2.5 EEG Acquisition and Data Preprocessing

We recorded EEG from 128 active electrodes using a Biosemi Active Two system (Biosemi, the Netherlands). Recordings were grounded using common mode sense and driven right leg electrodes (http://www.biosemi.com/faq/cms&drl.htm). We added 4 additional channels to the standard montage: two electrodes placed 1cm from the outer canthi of each eye, and two electrodes placed above and below the right eye. EEG was sampled at 512 Hz (DC-coupled with an anti-aliasing filter, −3dB at 102 Hz). Electrode offsets were kept within ± 40*μ*V.

We processed EEG data using EEGLab 13.4.4b (Delorme & Makeig, 2004) and Letswave 6 (http://nocions.webnode.com/letswave) running in MATLAB (The Mathworks). 50 Hz line noise was identified using Cleanline (Mullen, 2012) using a separate 1 Hz high-pass filtered dataset (EEGLab Basic FIR Filter New, zero-phase, finite impulse response, −6dB cutoff frequency 0.5 Hz, transition bandwidth 1 Hz). We subtracted this line noise from the unfiltered dataset (as recommended by Bigdely-Shamlo et al., 2015). We identified noisy channels by visual inspection (mean noisy channels by participant 0.6, median 0, range 0-3) and marked these for exclusion from average referencing and independent components analysis (ICA). We rereferenced the data to the average of the 128 scalp channels, and removed one extra channel (FCz) to correct for the rank deficiency caused by average referencing (as done by Feuerriegel et al., 2018). We processed a copy of this dataset in the same way, but additionally applied a 1 Hz high-pass filter (EEGLab Basic FIR Filter New, filter settings as above) to improve stationarity for the ICA. ICA was performed on the 1 Hz high-pass filtered dataset (RunICA extended algorithm, Jung et al., 2000). We then copied the independent component information to the unfiltered dataset (as recommended by Viola et al., 2010). We identified and removed independent components generated by blinks and saccades according to guidelines in Chaumon et al. (2015). After ICA we interpolated any bad channels (max bad channels within a dataset = 3) and FCz from the cleaned data (spherical spline interpolation).

### 2.6 Frequency Domain Data Processing

Following preprocessing, we epoched EEG data from 0-1000ms relative to each surprising oddball cycle onset, and from the onset of each expected oddball immediately preceding a surprising oddball. This ensured an equal number of epochs for each oddball type. The one-second epoch length was an exact multiple of the 166.6ms base rate stimulus cycle duration. For each oddball type, we concatenated epochs to produce 42-second sequences (the so-called ‘false-sequencing’ approach, see Quek & Rossion, 2017). Concatenated sequences contained periodic responses to oddball faces at a rate of 1 Hz (determined by the one-second epoch length) and periodic responses to base rate faces at 6 Hz. To avoid periodic signals from epoch edge artefacts, we adjusted the amplitude of the first sample in each epoch to match the amplitude of the last sample in the preceding epoch for each electrode. We then imported the resulting sequences into Letswave 6, high-pass filtered at 0.1 Hz (Butterworth 4^th^ order filter) and converted signals to the frequency domain using fast Fourier transforms (FFT; frequency resolution 0.0238 Hz).

#### 2.6.1 Z-Score Conversion

For Z-score conversion we first averaged Fourier amplitude spectra (also averaged across all channels and concatenated sequence types) across all participants, and then created z-scores for each frequency bin relative to the amplitudes of the 20 surrounding bins (excluding adjacent bins and the single bins with the highest and lowest amplitudes, see Retter & Rossion, 2016). We then assessed Z-scores resulting from the concatenated sequences at the exact frequency bin of each harmonic for oddball harmonics (multiples of 1 Hz) and base rate harmonics (multiples of 6 Hz) separately. We included harmonics in further analyses if the Z-score for that harmonic exceeded 3.1 (p<0.001, one-tailed, i.e., signal > noise) as done in previous studies (Jacques et al., 2016; Quek & Rossion, 2017). After the lowest frequency base rate or oddball harmonic was identified for inclusion in analyses, if any subsequently tested harmonic was not statistically significant at p<.001 then higher harmonics were not considered for further analyses. Z-scores of oddball harmonics were statistically significant from the 2^nd^ until the 7^th^ harmonic (i.e., 2, 3, 4, 5, 7, 8, 9 Hz, excluding the base rate of 6 Hz). Z-scores of base rate harmonics (6 Hz and higher multiples) were statistically significant until the 9^th^ harmonic (i.e., 6, 12, 18, 24, 30, 36, 42, 48, & 54 Hz).

#### 2.6.2 Summing Harmonics

To take into account noise variations across the amplitude spectrum, we baseline-corrected the frequency amplitude spectra for each channel separately using the 20 surrounding bins (excluding adjacent bins and single bins with highest and lowest amplitudes). We then summed the baseline-subtracted amplitudes at the exact frequencies of included significant harmonics, for oddball and base rate harmonics separately. To reduce the number of comparisons in statistical analyses, we averaged sums of harmonics across the expected Identity Plus Size Change oddballs in each sequence type for each participant. This averaging was done after sums-of-harmonics had been calculated for the expected oddballs in each sequence type separately, so that there was not a higher signal-to-noise ratio resulting from including more epochs for FFTs in the Identity Plus Size Change condition.

#### 2.6.3 Region of Interest (ROI) Definitions

We defined two ROIs for analyses: a right occipitotemporal ROI (PO8/10/12, P8/10) and a medial occipital ROI (Oz/1/2, POOz/5/6, OIz, POI1/2). The right occipitotemporal ROI is based on electrodes at which face identity oddball responses are typically largest (Liu-Shuang et al., 2014, 2016; Dzhelyova & Rossion, 2014; Xu et al., 2017). The medial occipital ROI was defined as a grid of electrodes at which base rate and oddball responses were largest when averaged across oddball types. For ROI analyses we averaged sums of baseline-subtracted harmonic amplitudes across electrodes within each ROI.

### 2.7 Task Performance Analyses

For analyses of task performance we calculated mean accuracies and response times for each sequence type, as well as sums of harmonics within each ROI, using 20% trimmed means and two-tailed 95% confidence intervals derived using the percentile bootstrap method (10,000 bootstrap samples; Efron & Tibshirani, 1994; Wilcox, 2012). We tested for differences in accuracy and reaction time across sequences using percentile bootstrapping of the between-sequence difference scores.

### 2.8 ROI Sums of Harmonics Analyses

For analyses of oddball and base rate harmonics at preselected ROIs, we conducted 2 × 2 × 2 repeated measures ANOVAs with the factors *Identity* (face identity change/identity repetition relative to base faces), *Size* (20% larger/same size as base faces) and *ROI* (medial occipital/right occipitotemporal). As we aimed to test for face identity repetition effects, we were interested specifically in the main effect of identity and interactions of identity × size, identity × ROI and identity × size × ROI, and so limited our analyses to these pre-specified effects (as opposed to assessing all main effects and interactions as done in exploratory analyses; see Cramer et al., 2014). Greenhouse-Geisser corrections were applied where appropriate. We corrected p-values for multiple comparisons across selected F tests (i.e. the main effects and interactions listed above) using the Holm-Bonferroni method (Holm, 1979) using the calculator by Gaetano (2013).

### 2.9 Mass Univariate Analyses Sums of Harmonics Analyses

We also tested for oddball-evoked frequency domain responses outside of predefined ROIs for Identity Change and Image Repetition oddballs. This was done to characterise the scalp topography of expectation effects (i.e., Identity Repetition oddball responses) and repetition suppression (i.e., Identity Change oddball responses). Quantifying each effect across the scalp is important to better understand each phenomenon, as both expectation and repetition effects have been found in frontal regions when recording fMRI BOLD responses (e.g., Grotheer & Kovacs, 2015; Wig et al., 2005, 2009) which would be missed by targeted ROI analyses. We tested for summed oddball harmonics that were above zero using a one-sample cluster-based permutation test based on the cluster mass statistic (Bullmore et al., 1999; Maris & Oostenveld, 2007) with a family-wise alpha level of 0.05. We used cluster-based permutation tests as they are sensitive to detect broadly-distributed effects while controlling the weak family-wise error rate (Maris & Oostenveld, 2007; Groppe et al., 2011). All 128 scalp electrodes were included (128 total comparisons). We performed Yuen’s single-sample t-test (Yuen, 1974; Wilcox, 2012) for each electrode using the original data and 10,000 random within-subject permutations. For each permutation, all t statistics corresponding to uncorrected p-values of <0.05 were formed into clusters with any neighbouring such t-scores. We defined spatial neighbours using the spatial neighbourhood matrix supplied in the LIMO toolbox (Pernet et al., 2011). The sum of the t statistics in each cluster is the ‘mass’ of that cluster; we used the most extreme cluster masses in each of the 10,000 permutation tests to estimate the null hypothesis distribution. We then derived cluster-level p-values by calculating the percentile rankings of cluster masses from the observed data relative to the null distribution. The p-value of each cluster was assigned to all members of the cluster; electrodes not included within a cluster were given a p-value of 1. We also added a minimum cluster size constraint so that statistically significant clusters were required to include 3 or more electrodes. This constraint was added as we found that cluster-based permutation testing with this constraint controlled the weak family-wise error rate at alpha = 0.05 using random samples of normally-distributed data with the same number of tests and the same sample size, and also when using random partitions of data from Experiment 1 (estimated weak family-wise error rates between 0.04-0.05). For the permutation tests we calculated corrected p-values according to the conservative method in Phipson and Smyth (2010).

## 3. Experiment 1 Results

### 3.1 Behavioural Task Performance

Target detection accuracy was near ceiling and did not differ across sequence type (group mean accuracy ranged between 98-99% across sequence types, overall accuracy 98.7% ± 2.4%, all p-values >0.68). Mean response time for correct responses was 438 ± 41ms (group mean RTs ranged from 437-440ms across sequence types). Response times did not significantly differ across sequence types (all p-values >0.35, 95% CIs: sequence type 1-2 = [-10.5, 2.8]; sequence type 1-3 = [-7.6, 3.7]; sequence type 2-3 = [-4.0, 7.1]).

### 3.2 Frequency Domain Results

#### 3.2.3 Sums of Harmonics ROI Analyses

Grand-averaged Fourier amplitude spectra for each oddball type at medial occipital and right occipitotemporal ROIs are displayed in Figure 3A. Head plots of summed baseline-subtracted oddball and base rate harmonics at each electrode are displayed in Figure 3B. All oddball types, except for Image Repetition oddballs, showed clear responses at oddball presentation frequencies and higher harmonics over posterior electrodes.

**Figure 3.**
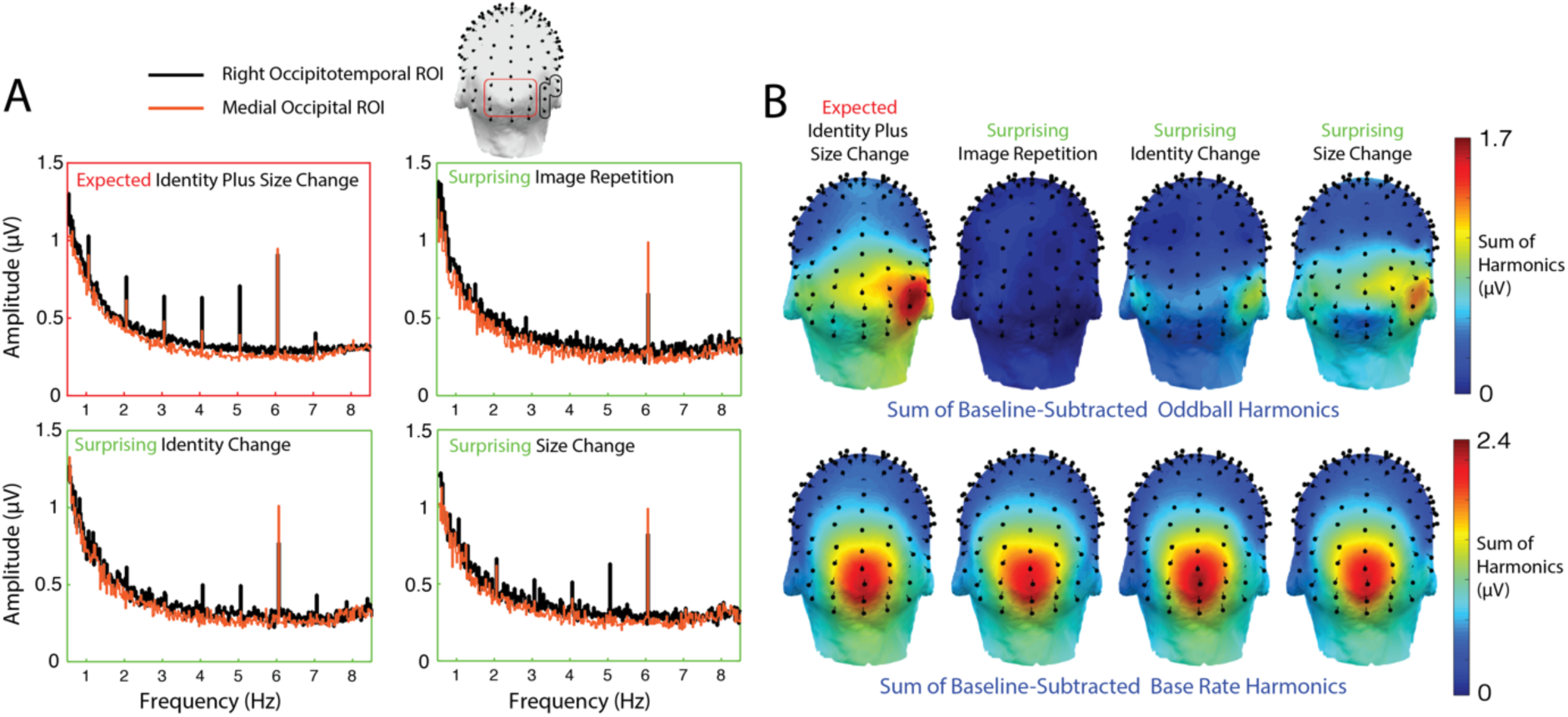
Oddball and base rate frequency domain responses in Experiment 1. A) Grand-averaged Fourier amplitude spectra for each oddball type at medial occipital and right occipitotemporal ROIs. B) Head plots of sums of baseline-subtracted oddball and base rate harmonics for each oddball type.

Grand averages of summed baseline-subtracted oddball harmonics for each oddball type at medial occipital and right occipitotemporal ROIs are displayed in Figure 4A. The response evoked by Image Repetition oddballs (i.e., surprising exact repetitions of the base rate face) was not significantly different from zero in both ROIs (medial occipital = 0.03, 95% CI = [-0.12, 0.19]; right occipitotemporal = 0.05, 95% CI = [-0.14, 0.25]). In contrast, there were significant oddball-selective response to Identity Change oddballs (indexing face identity change effects; medial occipital = 0.46, 95% CI = [0.26, 0.65]; right occipitotemporal = 0.80, 95% CI = [0.47, 1.14]), Size Change oddballs (indexing low-level stimulus changes, medial occipital = 0.60; 95% CI = [0.40, 0.89]; right occipitotemporal = 1.06; 95% CI = [0.81, 1.37]) and Expected Identity Plus Size Change oddballs (indexing low-level and identity changes, medial occipital = 0.90; 95% CI = [0.70, 0.17]; right occipitotemporal = 1.45; 95% CI = [1.11, 1.82]).

**Figure 4.**
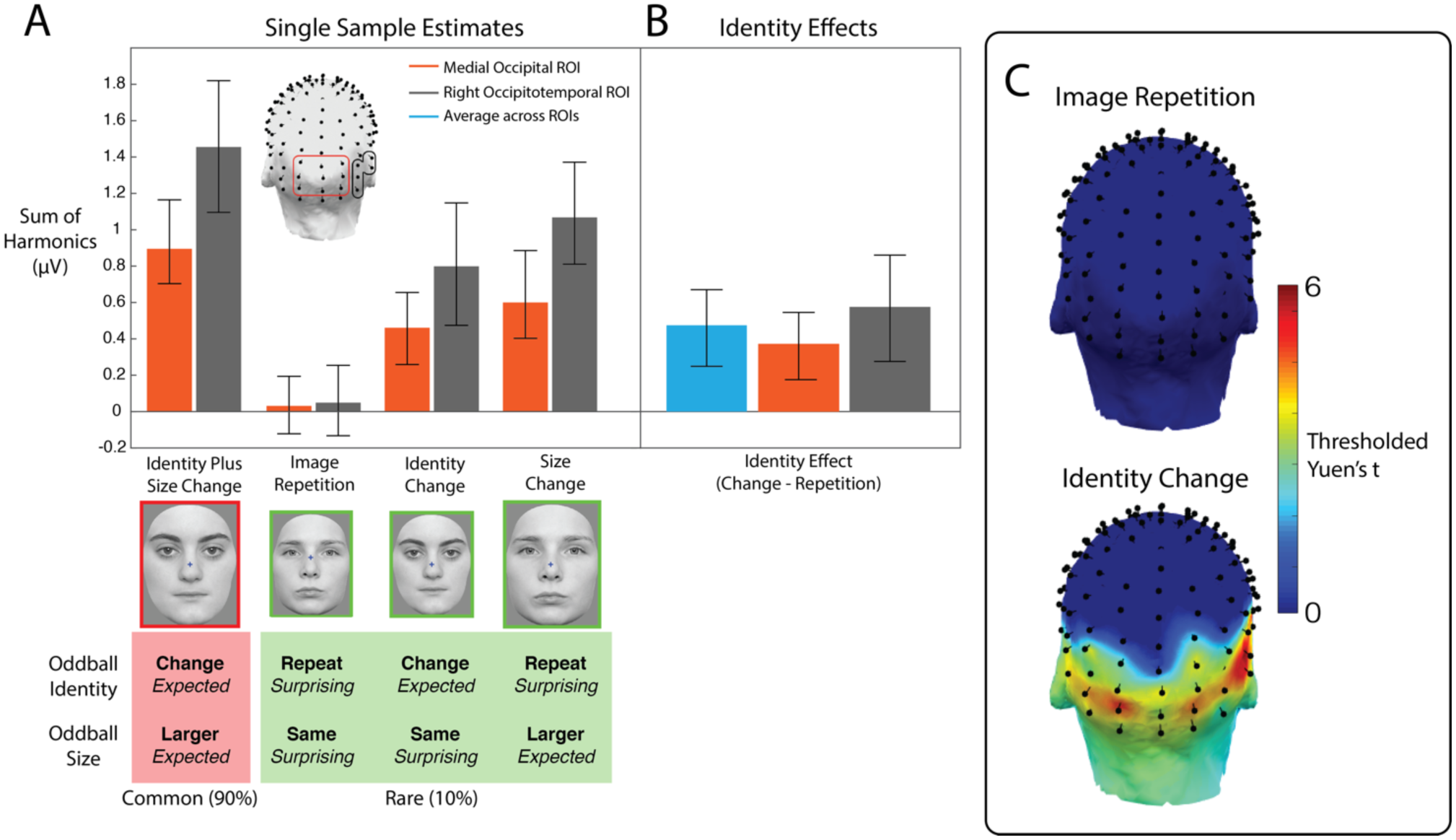
Results of Experiment 1. A) Sums of baseline-subtracted oddball harmonics for each oddball type at medial occipital and right occipitotemporal ROIs. Summed harmonics are multiples of the 1 Hz oddball frequency within the concatenated sequence (2-8 Hz, excluding the base rate of 6 Hz). Descriptions of each oddball’s face identity and size relative to base rate faces are written in bold text. The expectation status of each oddball stimulus characteristic is provided in italicised text. Error bars depict 95% confidence intervals; where these exclude zero this is indicative of statistically significant periodic oddball responses. B) Differences in oddball-evoked responses between identity change and identity repetition oddballs, averaged over size conditions. C) Results of mass univariate single sample tests for Image Repetition oddballs (i.e., indexing expectation effects) and Identity Change oddballs (i.e., indexing face identity repetition effects). Yuen’s t statistics are thresholded for statistical significance using cluster-based permutation testing.

To test for face identity repetition effects (i.e., smaller responses for oddballs of the same face identity to base faces), we conducted a 2 × 2 × 2 repeated measures ANOVA with the factors *Identity* (face identity change/identity repetition), *Size* (20% larger/same size as base faces) and *ROI* (medial occipital/right occipitotemporal). Summed oddball harmonics were larger for oddball faces with an identity change relative to base faces (main effect of identity, F(1,21) = 19.35, corrected p = 0.001, see Figure 4B). No other preselected main effects or interactions were statistically significant (all uncorrected p-values >0.05).

We also conducted a 2 × 2 × 2 repeated measures ANOVA with the same factors to test for differences in summed base rate harmonics (i.e., general differences in visual evoked responses across sequences) as a control measure. No preselected main effects or interactions were significant (all uncorrected p-values >0.05).

#### 3.2.4 Sums of Harmonics Mass Univariate Analyses

To test for expectation and repetition effects outside of predefined ROIs, mass-univariate single sample tests were performed for sums of baseline-subtracted oddball harmonics evoked by Identity Repetition oddballs (indexing expectation/surprise effects) and Identity Change oddballs (indexing face identity repetition effects; results displayed in Figure 4C). For the Identity Repetition oddballs (indexing expectation effects) there were no significant clusters of oddball responses. For Identity Change oddballs (indexing repetition effects) there was a cluster of responses spanning bilateral posterior electrodes (cluster mass = 100.67, critical cluster mass = 2.03, p <0.001).

## 4. Experiment 1 Interim Discussion

In Experiment 1 we showed that expectations of FPVS oddball stimuli do not *generate* an oddball response independently of stimulus-evoked responses. Point estimates and 95% confidence intervals for Image Repetition oddball responses in Figure 4A (which capture expectation/surprise responses that cannot be accounted for by face repetition effects) indicate that expectation or anticipation processes alone does not generate oddball responses in our FPVS-EEG design. Given that in previous FPVS-EEG studies, expectation or anticipation effects are even reduced by other factors (i.e., random size variation at the base rate, and occurrence of different face identities at oddballs, e.g., Liu-Shuang et al., 2014; 2016; Dzhelyova & Rossion, 2014; Dzhelyova et al., 2017; Xu et al., 2017), these observations indicate that oddball responses in FPVS-EEG designs are not due to expectation or anticipation effects. Instead, they appear to primarily index neural responses driven by the perceived periodic stimulus changes relative to base rate stimuli.

## 5. Experiment 2

A complementary approach to Experiment 1 is to ask whether oddball-evoked responses can be *modulated* by expectation and surprise (see Quek & Rossion, 2017). We tested this possibility in Experiment 2 by presenting oddball faces that were 20% larger than base rate faces (found to generate measurable oddball responses in Experiment 1) in expected (common) and surprising (rare) contexts. We presented oddballs that were the same face identity as base faces (identity repetitions) and different faces (identity change oddballs) in these contexts, and quantified oddball-evoked responses in the frequency domain using the ‘false sequencing’ approach as in Experiment 1. This approach allowed us to assess the relative size of expectation effects for immediately repeated and unrepeated stimulus identities. We did this to test the hypothesis that stimulus expectation effects are diminished or abolished when stimuli are repeated, based on patterns of results from existing fMRI studies (e.g., Kovacs et al., 2012; Larsson & Smith, 2012; Symonds et al., 2016). It is important to note that repetition as described here refers to *immediate* repetition (i.e., stimuli that are repeated without any intervening stimuli), as the expectation effects in Experiment 2 could also be conceptualised as *delayed* repetition effects (i.e., driven by frequent presentations of the expected stimuli which are separated by several intervening stimuli; see Vinken & Vogels, 2017). If expectation effects are reduced for repeated stimuli, this may also explain why we did not observe surprise effects for Image Repetition oddballs in Experiment 1, as surprise responses would have been heavily suppressed by massed repetition of the base rate face.

We expected to find larger frequency domain expected/surprising differences for unrepeated (compared to repeated) face identity oddballs, based on previous findings from non-oddball designs that found larger expectation effects for unrepeated stimuli (e.g., Todorovic & de Lange, 2012; Symonds et al., 2016). We also extracted time-domain waveforms evoked by oddball stimuli to characterize visual mismatch responses in the context of FPVS-EEG designs.

### 5.1 Participants

Eighteen people participated in Experiment 2 (4 males, age range 18-33 years, mean age 23 ± 4.3 years). All participants had normal or corrected-to-normal vision, no history of psychiatric or neurological disorders or substance abuse and were right-handed as assessed by the Flinders Handedness Survey (Nicholls et al., 2013). All participants showed unimpaired facial recognition ability as measured by the electronic version of the Benton Facial Recognition Test (all global accuracy scores > 39/54 cutoff defined in Rossion & Michel, 2018). Experiment 2 was approved by the Human Research Ethics committee of the University of South Australia.

### 5.2 Stimuli

We took eighteen frontal images of faces (8 males, 10 females, neutral expression) from Laguesse and Rossion (2013) and processed these as in Experiment 1. Stimuli subtended 6.7° × 9.1° visual angle at a distance of 60cm. We created a separate, larger set of faces by scaling the resulting face images by 120%.

### 5.3 Procedure

Participants sat in a well-lit room 60cm in front of an LED monitor (refresh rate 60 Hz). We presented stimuli using PsychToolbox V.3.0.11 (Kleiner et al., 2007) running in MATLAB (Mathworks) against a grey background. Stimulation sequence structure and experimental task were identical to Experiment 1, except that participants responded using a one-button response box connected directly to the EEG amplifier.

We presented 4 types of oddball stimuli in Experiment 2 (displayed in Figure 5). All oddballs were 20% larger than base rate faces. Oddballs were either a different identity relative to base rate faces (Identity Change) or the same identity (Identity Repetition). The Identity Change oddball was of a single face identity, which did not change throughout a given sequence. Identity Change and Identity Repetition oddballs were presented in two different sequence types within the experiment. In the Identity Change Common sequences the Identity Change oddballs made up 90% of all oddballs (i.e., an identity change at the oddball frequency is expected) and Identity Repetition oddballs appeared as 10% of oddballs (i.e., an identity repetition at the oddball frequency is surprising). To clarify, identity repeats could be expected for base rate faces, but it was surprising to see an identity repetition of the base rate faces at the time of oddball stimulus appearance. We reversed these stimulus probabilities for the Identity Change Rare sequences.

**Figure 5.**
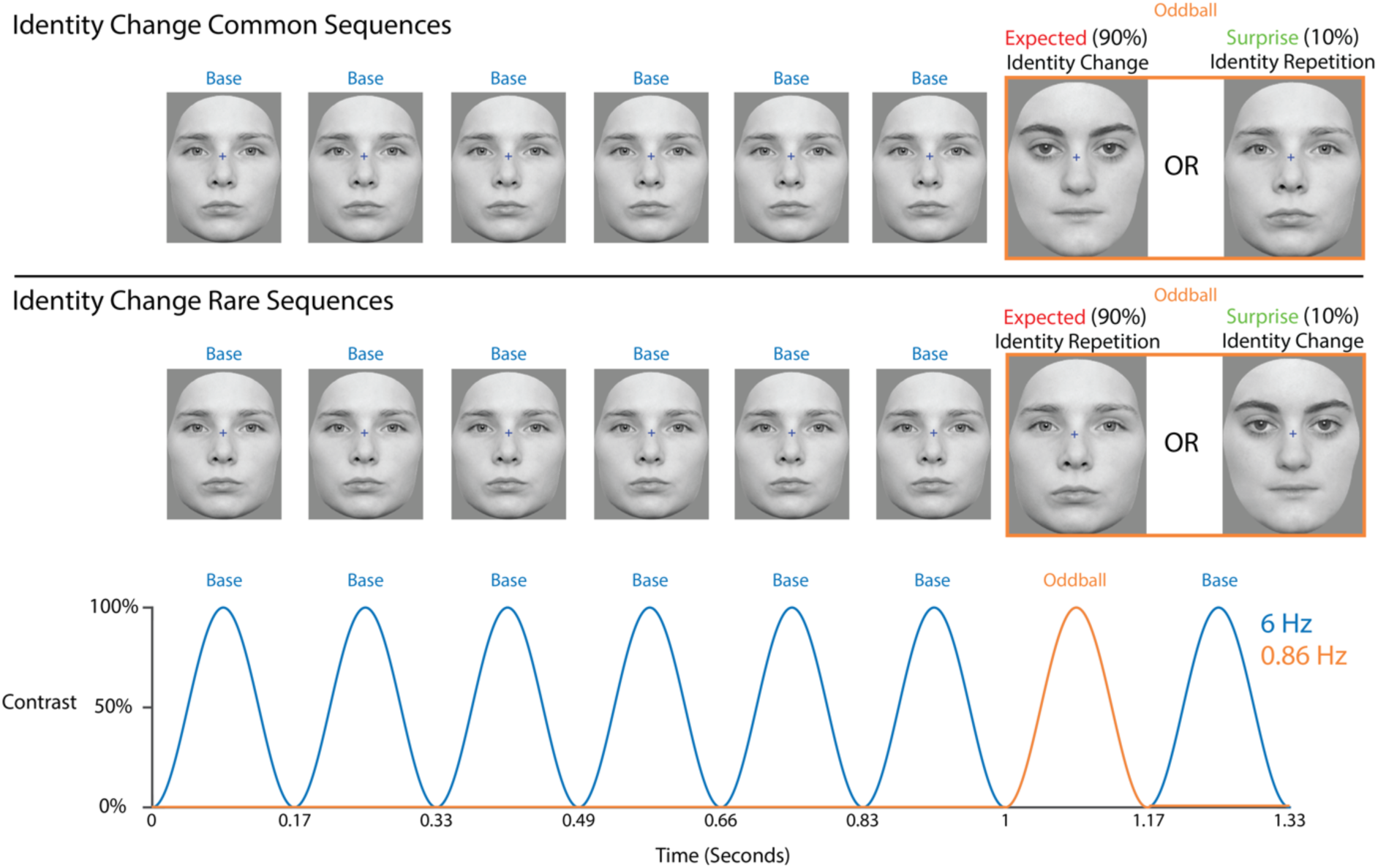
Experiment 2 stimulation sequence diagram. Faces were presented using sinusoidal contrast modulation at a rate of 6 cycles per second (i.e., 6 Hz). Oddball faces (outlined in orange) appeared after every 6 base faces at a rate of 0.857Hz, and were always 20% larger than the base rate faces Within each sequence there were two oddball stimulus types: a different face identity to base rate faces (Identity Change oddball) and the same face identity as base rate faces (Identity Repetition oddball). The Identity Change oddball was of a single face identity which did not change throughout the sequence. On a given trial, one of these oddball stimulus type was expected (90% appearance probability) and the other was surprising (10% appearance probability).

Within a sequence we presented pairs of either male or female faces as base and oddball stimuli (4 sequences male faces, 5 sequences female faces for each sequence type). Each pair of face images was presented in one sequence each for both sequence types. There were nine sequences of each sequence type, each containing seven surprising oddball instances for a total of 63 surprising oddballs per sequence type across the full experiment. We counterbalanced face identities presented as base faces and Identity Change oddballs across participants. Total testing duration was 27 minutes.

### 5.4 Task Performance Analyses

We estimated mean accuracies and response times for Identity Change Common and Identity Change Rare sequences as in Experiment 1, using 20% trimmed means and 95% confidence intervals using the percentile bootstrap (10,000 bootstrap samples).

### 5.5 EEG Acquisition and Data Processing

We performed EEG data acquisition and preprocessing as in Experiment 1, with the following changes: we added 8 rather than 4 additional channels to the standard montage, including two electrodes placed 1cm from the outer canthi of each eye, four electrodes placed above and below each eye, and two electrodes placed on the left and right mastoids. The mean number of excessively noisy electrodes per participant was 1.3 (median 1, range 0-6).

### 5.6 Frequency Domain Statistical Analyses

For frequency domain analyses we performed frequency domain preprocessing, z-scoring, baseline-subtraction and summing of harmonics as in Experiment 1, for expected and surprising Identity Change and Identity Repetition oddball stimuli, with the following differences. In Experiment 2 concatenated sequences were 63 seconds in duration (FFT frequency resolution 0.0159 Hz). We assessed differences in sums of oddball and base rate harmonics by stimulus probability (expected/surprising) for Identity Change and Identity Repetition oddballs using mass univariate paired-samples Yuen’s t tests with cluster-based multiple comparisons corrections as described in Experiment 1. We also tested for a repetition by stimulus probability interaction effect by comparing expected/surprising within-subject difference scores for Identity Change and Identity Repetition oddballs.

### 5.7 Time Domain Data Processing

For time domain analyses EEG data were high-pass filtered at 0.1 Hz (EEGLab Basic Finite Impulse Response Filter New, non causal zero-phase, −6dB cutoff frequency 0.05 Hz, transition bandwidth 0.1 Hz) and then low-pass filtered at 30 Hz (EEGLab Basic Finite Impulse Response Filter New, non causal zero-phase, −6dB cutoff frequency 33.75 Hz, transition bandwidth 7.5 Hz). As we were primarily interested in ERPs evoked by the oddballs (and not the base rate faces) data from each sequence were notch filtered at the 6 Hz base rate and higher harmonics (first 5 harmonics spanning 6-30 Hz; stopband width 0 Hz, slope cutoff width 0.05 Hz) to remove time-domain responses evoked by base rate stimuli (as done by Dzhelyova & Rossion, 2014; Rossion et al., 2015; Retter & Rossion, 2016).

We then epoched the resulting data from −166ms to 834ms time-locked to the start of i) each surprising oddball stimulus cycle, and ii) the immediately preceding expected oddball cycle. We baseline-corrected the resulting epochs using the prestimulus interval. Epochs containing ±100μV deviations from baseline in any of the 128 scalp channels were excluded from analyses (minimum of 60 epochs retained per oddball type per participant).

### 5.8 Time Domain Statistical Analyses

We compared time domain waveforms evoked by expected and surprising oddballs for Identity Change and Identity Repetition stimulus types separately, using mass univariate paired-samples Yuen’s t tests performed in the LIMO EEG toolbox V1.4 (Pernet et al., 2011). All time points between −166 and 834ms at all 128 scalp electrodes were included in each test. We performed corrections for multiple comparisons using spatiotemporal cluster corrections with a cluster inclusion significance threshold of 0.05 and 1000 bootstrap samples to estimate the null distribution. As for Experiment 1 we defined which channels were spatial neighbours using the 128-channel Biosemi channel neighbourhood matrix in the LIMO EEG toolbox (Pernet et al., 2011). Adjacent time points were considered as temporal neighbours.

To determine whether stimulus repetition modulated expectation/surprise effects, we conducted mass univariate 2 × 2 repeated measures ANOVAs with the factors *stimulus probability* (expected/surprising) and *face identity* (identity change/identity repetition). We used F ratios as test statistics for mass univariate analyses in this case. Main effects in this ANOVA model were not assessed as they were not of interest in this study.

One potential confound in this experiment is that the expected oddball face image in each sequence is repeated more times compared to the surprising oddball in the same sequence. Because of this, expected oddballs may show stronger delayed (across-oddball) repetition effects (see Henson et al., 2004; Xiang & Brown, 1998). Delayed repetition refers to when stimuli (i.e., the oddballs) are repeated after a number of intervening stimuli (i.e., the base faces), and is associated with systematic changes in ERPs (Henson et al., 2004); single cell firing rates (Xiang & Brown, 1998) and BOLD signals (Henson et al., 2000; Sayres & Grill-Spector, 2006), with stronger effects after more repetitions of the same image. To evaluate this, we tested for changes in oddball-evoked waveforms by position in the sequences for expected Identity Change and Identity Repetition oddballs. These analyses were run to reveal whether time domain waveforms evoked by expected oddballs systematically changed over the course of the sequence (i.e., became more positive or negative with more oddball image repetitions). For these analyses, responses to *all* expected oddballs in each sequence (not only those preceding a surprising oddball) were epoched as above and included in analyses. We performed within-subject linear regressions on the amplitudes at each time point and electrode combination with presentation number of the expected oddball within a sequence (ranging from 1-66) as a predictor. Beta coefficients were estimated for each subject at each time point and electrode combination. At the group-level we performed single-samples Yuen’s t tests on the beta coefficients, to assess whether they differed from zero, using cluster-based multiple comparisons corrections as described above. Beta coefficients larger than zero indicate that waveforms evoked by oddballs presented later in the sequences were more positive than for oddballs presented early in the sequences. Conversely, beta coefficients smaller than zero indicate that amplitudes became more negative for the expected oddballs presented later in the sequences.

## 6. Experiment 2 Results

### 6.1 Task Performance

As in Experiment 1, target detection accuracy was near ceiling (mean accuracy ranging between 98-99% for both Identity Change Common and Identity Change Rare sequence types, overall accuracy 98.5% ± 1.6%). Mean response time for correct responses was 401.6 ± 27.2ms. Accuracies and response times did not significantly differ across sequence types (Identity Change Common – Identity Change Rare 95% CIs: accuracy [-0.48%, 1.67%], mean response time [-7.8ms, 3.3ms]).

### 6.2 Frequency Domain Results

Z-scores were calculated as in Experiment 1 by averaging across all participants, conditions and channels to identify harmonics to be included in further analyses. Oddball harmonics in the concatenated sequences were statistically significant until the 8^th^ harmonic (i.e., 1, 2, 3, 4, 5, 7, 8, 9 Hz, excluding the base rate of 6 Hz). At the group level, Z-scores of base rate harmonics (6 Hz and higher multiples) were statistically significant until the 9^th^ harmonic (i.e., 6, 12, 18, 24, 30, 36, 42, 48, & 54 Hz). Head plots of summed baseline-subtracted oddball and base rate harmonics at each electrode are displayed in Figure 6A and 6B.

**Figure 6.**
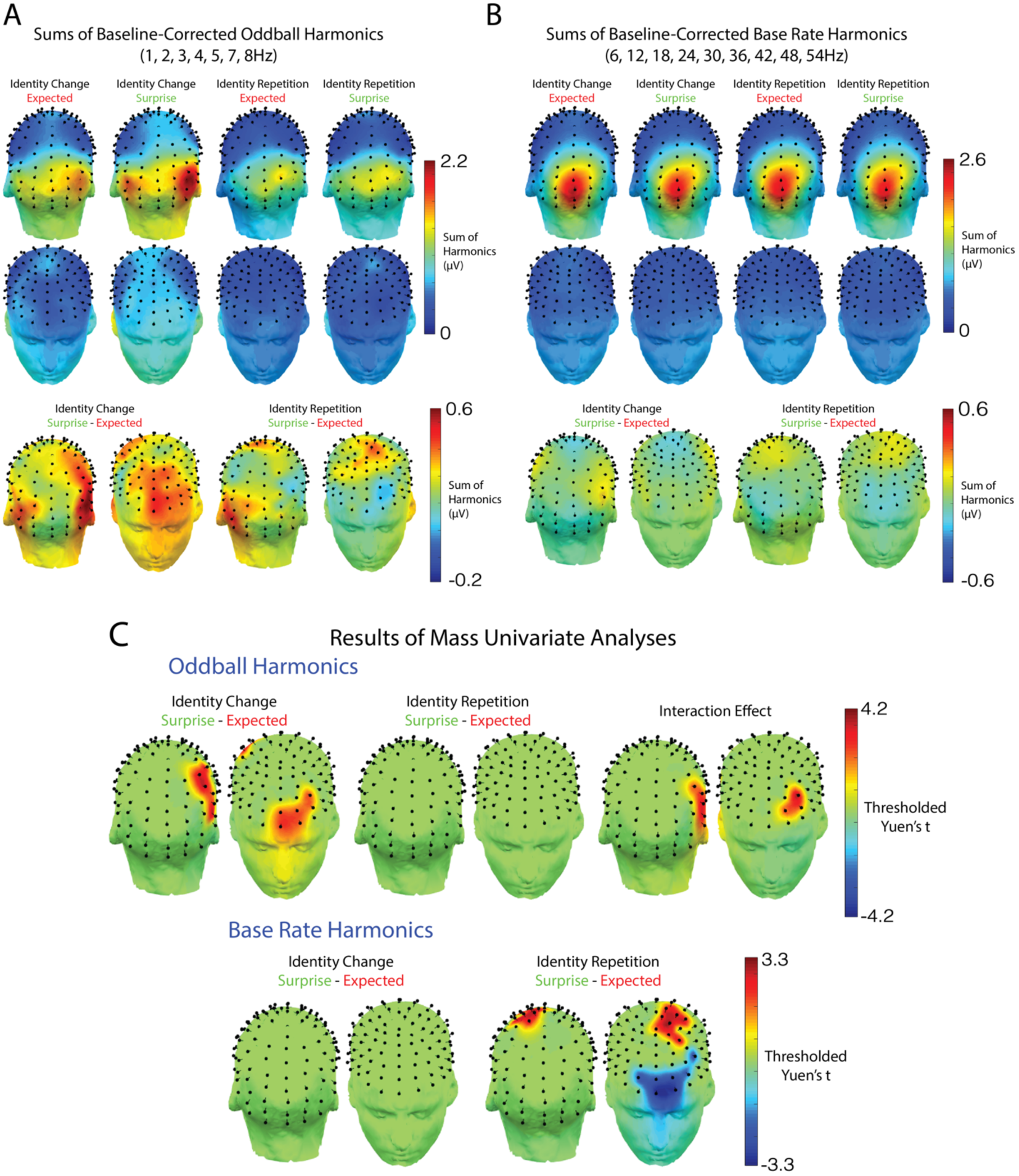
Frequency domain results for Experiment 2. A) Top and Middle Rows: Head plots of summed of baseline-subtracted oddball harmonics for each oddball type. Bottom Row: Conditional difference topographies (surprise-expected). B) Head plots of summed baseline-subtracted base rate harmonics for each oddball type, and (surprise-expected) conditional difference topographies. C) Results of mass univariate paired samples tests on summed baseline-subtracted oddball harmonics (top row) and base rate harmonics (bottom row). Yuen’s t statistics are thresholded for statistical significance using cluster-based permutation testing.

To test for expected/surprising differences in the sums of oddball and base rate harmonics, mass univariate tests with cluster-based multiple comparisons corrections were performed over all electrodes, for Identity Change and Identity Repetition oddballs separately. Mass univariate analyses of expected/surprising differences in summed oddball harmonics revealed two statistically-significant clusters at which responses were larger when evoked by surprising compared to expected Identity Change oddballs, displayed in Figure 6C: a right occipitotemporal cluster (cluster mass = 20.16, critical cluster mass = 3.90, p <0.001, electrodes CPP4h, CPP4, CCP4, CPP6h, P6, P8) and a left frontal cluster (cluster mass = 15.31, critical cluster mass = 3.90, p <0.001, electrodes Fpz, Afpz, AF3, Fp1, AFF5, AFF5h). There were no significant clusters of expected/surprising differences for Identity Repetition oddballs. Expected/surprising differences were larger for Identity Change compared to Identity Repetition oddballs in a right occipitotemporal cluster (cluster mass = 18.78, critical cluster mass = 4.26, p <0.001, electrodes CPP4, CPP6h, P6, TP8h, P8, TP8, P10) and a left frontal (cluster mass = 9.78, critical cluster mass = 4.26, p <0.001, electrodes AF3, AFF5, AFF5h).

Analyses of summed base rate harmonics did not reveal any significant stimulus expectation effect clusters for Identity Change oddballs (Figure 6C). For Identity Repetition oddballs there were two significant clusters: a left-lateralised central cluster at which responses were larger to surprising oddballs (cluster mass = 24.36, critical cluster mass = 12.09, p = 0.001, electrodes AF4, Fpz, AFpz, AF3, Fp1, AFF5, FFC5h) and a frontal cluster at which responses were larger to expected oddballs (cluster mass = 19.38, critical cluster mass = 12.09, p = 0.008, electrodes FCz, FFC1, FCC1h, FCC1, FFC3h, C1, C1h, CCP1h, C3).

### 6.3 Time Domain Results

Grand average time domain waveforms evoked by expected and surprising oddballs are shown in Figure 7A. We conducted mass univariate analyses of time domain waveforms evoked by expected and surprising oddballs for Identity Change and Identity Repetition oddballs. For Identity Change oddballs, time domain waveforms between 206-358ms at parieto-occipital electrodes were more negative when identity changes were surprising than when they were expected (cluster mass = 3738.24, critical cluster mass = 73.41, p = 0.001) as displayed in Figure 7B. No significant stimulus probability effect clusters were identified for Identity Repetition oddballs.

**Figure 7.**
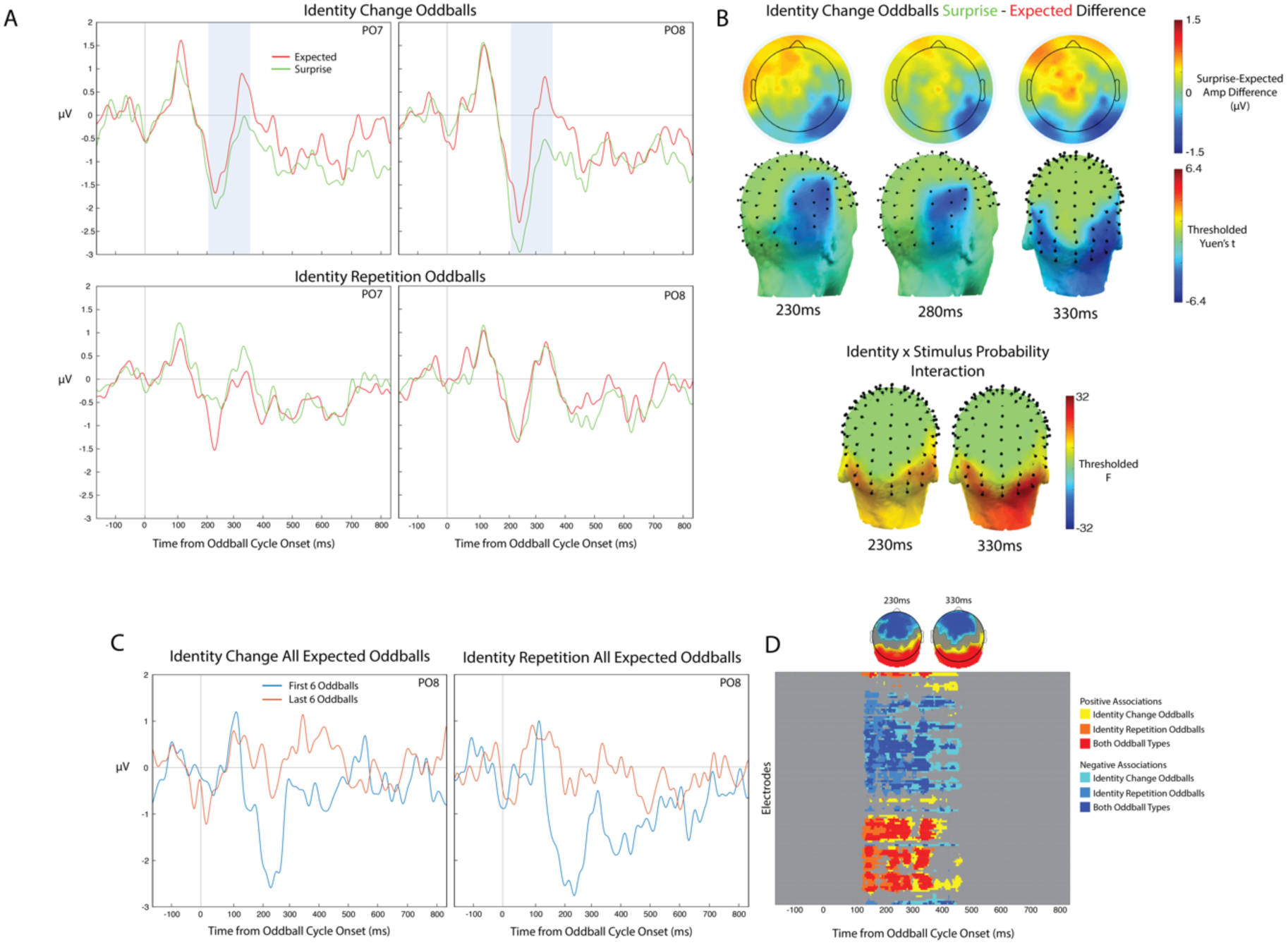
Experiment 2 time domain results. A) Grand average time domain waveforms evoked by each Identity Change oddballs (top row) and Identity repetition oddballs (bottom row). Blue shading denotes periods during which statistically significant stimulus probability (expected/surprise) effects were found using mass univariate analyses. B) Expected-surprise amplitude difference maps and head maps showing topographies of statistically significant effects at different latencies from oddball cycle onset. Yuen’s t and F statistics were thresholded for statistical significance using cluster-based permutation testing. C) Grand average time domain waveforms of the averages of the first and last 6 expected oddballs within a sequence for Identity Change and Identity Repetition oddballs. D) Group-level results of sequence position effect analyses for expected Identity Change and Identity Repetition oddballs. Each colour denotes the time point/electrode combinations at which a positive or negative effect of sequence position (positive or negative betas) were found, corrected for multiple comparisons using cluster-based permutation testing. Positive associations denote more positive amplitudes for oddballs presented later in the sequences (i.e., with increasing numbers of within-sequence image repetitions). Negative associations denote more negative amplitudes for oddballs presented later in the sequences.

Mass univariate 2 × 2 repeated measures ANOVAs were conducted on time domain data to test for differences in the magnitude of expected/surprising differences across Identity Change and Identity Repetition oddball types. Stimulus expectation effects were larger for Identity Change compared to Identity Repetition oddballs between 212-244ms (cluster mass = 3155.00, critical cluster mass = 2314.71, p = 0.001) and between 286-360ms (cluster mass = 5233.69, critical cluster mass = 2314.71, p = 0.001), largely overlapping with the spatiotemporal topography of stimulus expectation effects for Identity Change oddballs (see Figure 7B).

#### 6.3.1 Sequence Position Effect Analyses

As the expected oddball images were presented many more times within a sequence compared to surprising oddballs, delayed or across-oddball repetition effects may have contributed to the observed expectation effects. To assess this, we ran mass univariate linear regression analyses to test whether waveforms evoked by expected oddball images systematically changed over the course of the sequences (i.e., whether waveforms became more positive or negative with increasing numbers of within-sequence oddball image repetitions). These analyses were run for both Identity Change and Identity Repetition oddballs. To visualise changes in oddball-evoked time domain waveforms across the duration of each sequence, grand-average waveforms of averages of the first and last 6 expected oddballs are plotted in Figure 7C. These plots reveal large effects of sequence position for both Image Change and Image Repetition oddballs, which are of similar magnitudes. Statistically significant clusters with largely overlapping time ranges and topographies were found for both Identity Change and Identity Repetition expected oddballs (Figure 7D). Positive correlation clusters (indicating where waveforms became more positive as the sequence progressed) were found over posterior electrodes for Identity Change (136-460ms) and Identity Repetition (128-364ms) expected oddballs. Negative correlation clusters were also found over frontal electrodes (Identity Change 136-460ms, Identity Repetition 128-444ms). These effects show that time domain waveforms became more positive at posterior channels and more negative over frontal channels for oddballs presented later within each sequence, for both Identity Change and Identity Repetition oddballs.

## 7. Experiment 2 Interim Discussion

In Experiment 2 we adapted our FPVS design to separate immediate face identity repetition (i.e., whether the oddball face was a repeat of the preceding base rate face) and expectation (i.e., the frequency at which an oddball image was presented within a sequence). This allowed us to test for stimulus expectation-related *modulation* (rather than generation) of oddball stimulus-evoked responses, and whether such expectation-related modulations are suppressed for immediately repeated face identities. We found that stimulus expectations modulated the magnitude of frequency domain responses when oddballs were faces of a different identity to base faces (i.e., Face Identity Change oddballs). Surprising face identity oddballs evoked larger frequency-domain responses than when these identities were expected (i.e., presented often) at right occipitotemporal, right parietal and left frontal electrodes. We also observed more negative-going time domain waveforms to surprising oddballs between 206-358ms. However, we did not observe similar effects when oddballs were an identity repetition of the base rate faces, indicating again (i.e., as in experiment 1) that expectation effects in our designs were reduced or absent in the presence of face identity repetition.

Results of our control analyses indicated that the abovementioned expectation effects are at least partially distinct from effects of delayed (across-oddball) repetitions, which were more frequent for expected compared to surprising oddball stimuli. Time domain waveforms evoked by oddballs presented later in the sequences (i.e., after many oddball image repetitions) systematically differed from those presented earlier in the sequences. This was the case for both identity repetitions and identity changes relative to base rate stimuli, with similar effect magnitudes for both oddball types. However visual mismatch effects were only found when oddball stimuli included an identity change. The gradual changes in time domain waveforms as the sequences progressed instead appear to index a general habituation of responses with continuous stimulation.

## 8. General Discussion

Although it is understood that both repetition suppression and expectation contribute to visual mismatch responses in human EEG, classical visual oddball designs can provide little insight into whether these underlying mechanisms are additive or interactive. Here we addressed this outstanding question using a visual oddball design adapted from the fast periodic visual stimulation (FPVS) paradigm (Liu-Shuang et al., 2014, 2016; Dzhelyova & Rossion, 2014; Dzhelyova et al., 2017; Xu et al., 2017), in which effects of immediate stimulus repetition and expectation are separable. In Experiment 1, we tested whether violations of stimulus expectations are able to *generate* EEG responses that could not be accounted for by stimulus repetition; in Experiment 2 we characterised how expectation *modulates* stimulus-evoked responses, and whether the magnitude of modulation differed for immediately repeated and unrepeated stimuli. The critical finding we report here is that expectation does indeed modulate stimulus-evoked responses, leading to visual mismatch responses, however such expectation-related modulations are reduced or absent for immediately repeated stimuli (i.e., face image or identity repetitions). Moreover, expectation violations alone do not appear to *generate* measurable EEG responses when the surprising stimulus was a repetition of an image seen immediately beforehand. Our results show that repetition suppression can reduce perceptual expectation effects on stimulus-evoked responses in oddball designs, contrary to the view that stimulus expectations modulate repetition suppression (Summerfield et al., 2008; for discussion see Grotheer & Kovacs, 2016; Feuerriegel et al., 2018).

### 8.1. Stimulus Repetition Inhibits Expectation Effects

In the data presented here, expectation effects resembling visual mismatch responses were reduced for repeated stimuli, and were only evident for stimuli that differed to the image seen immediately beforehand (i.e., unrepeated stimuli). That is, we found no evidence for visual mismatch-like expectation effects for critical stimuli (i.e., oddballs) that were the same image (Experiment 1) or identity (Experiment 2) as the faces preceding that stimulus (i.e., the base rate faces).

In Experiment 1 we found that violations of participants’ stimulus expectations did not *generate* a substantial EEG signal independently of stimulus-evoked responses. That is, there was no evidence of a vMMN-like response to unexpected repetitions of the base rate face. This finding complements those of Quek and Rossion (2017), who used a FPVS design with highly variable images to manipulate participants’ expectations for certain stimulus categories (e.g., faces amongst objects), but did not find category-level expectation or anticipation responses. Taken together, these findings suggest that even when participants can form very strong image-specific expectations (e.g., for a particular oddball face image), measurable expectation effects do not arise unless the oddball stimulus itself is changed relative to the image seen immediately beforehand. Results from Experiment 2 further support this conclusion. Here we characterised a vMMN effect driven by *modulation* of stimulus evoked responses, yet this effect was only evident for face oddballs that were of a different identity to the immediately preceding image (i.e., non-repetitions). As mentioned above (interim discussion of Experiment 1), given that in previous FPVS-EEG studies, expectation or anticipation effects are even reduced by other factors (i.e., random size variation at the base rate, and occurrence of different face identities at oddballs, e.g., Liu-Shuang et al., 2014; 2016; Dzhelyova & Rossion, 2014; Dzhelyova et al., 2017; Xu et al., 2017), these observation indicate that oddball responses in FPVS-EEG designs are not due to expectation or anticipation effects. Instead, they appear to primarily index neural responses driven by the perceived periodic stimulus changes relative to base rate stimuli.

The finding that expectation effects in visual oddball designs are larger in the presence of stimulus change (i.e., for unrepeated stimuli), is consistent with a substantial body of evidence from non-oddball designs. Previous studies have shown that response differences to expected/surprising stimuli are larger for unrepeated compared to repeated stimuli (Amado et al., 2016; Waconge et al., 2011; Todorovic & de Lange, 2012; Kovacs et al., 2012; Larsson & Smith, 2012; Symonds et al.,2016). That stimulus repetition appears to suppress expectation effects suggests that there are at least two interacting mechanisms which underlie visual oddball effects, similar to models proposed to account for auditory mismatch responses (Costa-Faidella, et al., 2011; Mittag et al., 2016). The two mechanisms described in these models broadly correspond to “local” and “global” predictions as defined in Waconge et al. (2011): Local predictions are based on recent stimulus exposure (operationalised as stimulus repetition). Global predictions are based on the contextual probabilities of different events or stimulus sequences, operationalised as the presentation frequency (common/rare) of different oddball stimuli in our experiments. Critically, our findings extend these models by showing that such hierarchies can operate in fast periodic visual oddball designs, and that when local predictions are fulfilled (e.g., through stimulus repetition), violations of global probability rules lead to smaller or absent EEG waveform modulations than for unrepeated stimuli.

What are the likely cortical sources of the visual mismatch responses we observed in Experiment 2? The topographies of the frequency domain (Figure 6C) and time domain (Figure 7B) mismatch effects suggest contributions from electrical dipoles in both visual and frontal regions. The earlier phase of the time domain visual mismatch response (220-280ms in Figure 7B) appears to reflect a posterior negativity to surprising stimuli, accompanied by the negative dipole of a frontal positivity to surprising stimuli (for similar results see Dambacher et al., 2009; Symonds et al., 2017). The later part of the response (~330ms in Figure 7B) appears to instead be generated from bilateral posterior sources. Our observations are congruent with expectation effects found over frontal electrodes in EEG (Feuerriegel et al., 2018; Hall et al., in press), and in BOLD signals in frontal areas such as inferior frontal and middle frontal gyri (Grotheer & Kovacs, 2015; Amado et al., 2016) and dorsolateral prefrontal cortex (den Ouden et al., 2009; Rahnev et al., 2011), as well as in ventral temporal regions when presenting face stimuli (e.g., Egner et al., 2010; Grotheer & Kovacs, 2015). Our results also provide further evidence that expectation effects underlying VMRs are not restricted to visual areas, as is commonly assumed in existing studies of visual mismatch responses. Future studies using visual oddball designs could evaluate effects across the entire scalp (i.e., not only at posterior electrodes) in order to better capture multiple sources of expectation effects (see Hall et al., 2018).

### 8.2. Implications for Models of Repetition Suppression and Perceptual Expectations

The models of mismatch responses described in the previous section are also similar to more general multistage models of repetition effects, which describe qualitatively different repetition and expectation effects across sensory hierarchies (Grimm et al., 2016; Grotheer & Kovacs, 2016; Henson, 2016). Our findings also extend these models to include a mechanism wherein stimulus repetition effects modulate the expression of expectation or surprise effects. This might occur via reductions in stimulus salience with repetition (Kaliukhovich & Vogels, 2016) corresponding to decreases in the precision of predictions and reduced expectation violation responses within predictive coding frameworks (Feldman & Friston, 2010; Auksztulewicz & Friston, 2016). Mechanistically, this may occur due to stimulus repetition-induced alterations of local excitatory/inhibitory neural circuit dynamics, which can lead to imbalances of lateral inhibition among competing feature-selective neurons (see Kaliukhovich & Vogels, 2016 for evidence supporting this circuit model). In this case lateral inhibition is skewed toward suppressing responses of excitatory neurons tuned to features of the repeated stimulus, further reducing their responses. Such repetition-induced changes in circuit dynamics could minimise later influences of expectation or attention-related modulations, that presumably operate on the same stimulus-selective neurons through feedback or lateral inhibitory connections (Auksztulewicz & Friston, 2016; Reynolds & Heeger, 2009).

Another possibility is that expectation effects as observed in Experiment 2 are caused by gain modulations of stimulus-evoked responses, resulting in amplified responses for surprising compared to expected stimuli (e.g., Larsson & Smith, 2012). In our experiments, response gain modulations may have been reduced, and therefore not detected, when stimulus-evoked responses were already heavily suppressed by massed repetition of the base rate face identity. This gain modulation mechanism would be distinct from processes generating mismatch responses independently of stimulus-evoked signal magnitude (for discussion of the arguments for and against such processes see Stefanics et al., 2014; May & Tiitinen, 2010). To distinguish between gain modulation effects and ‘endogenous’ mismatch processes future experiments could present high and low contrast stimuli to assess whether surprise effects scale with evoked response magnitude (for examples of this manipulation in attention research see Lee & Maunsell, 2009).

### 8.3 Stimulus Repetition Effects Persist When Repetitions are Surprising

In Experiment 1 we also observed face identity repetition suppression (i.e., smaller evoked responses for oddballs that were repetitions of the base face identity), despite identity repetitions being highly unlikely (i.e., surprising). This provides further evidence against the hypothesis that repetition suppression simply reflects perceptual expectations, as described in predictive coding models of repetition effects (e.g., Summerfield et al., 2008; Auksztulewicz & Friston, 2016). In these models, immediate repetition effects arise due to a default expectation that the most recently viewed stimulus will appear again. Repeated stimuli are hypothesised to elicit suppressed responses because they fulfil this default expectation. However, according to predictive coding accounts repeated stimuli will evoke *enhanced* responses (i.e., larger than to unrepeated stimuli) when stimulus repetitions are made unlikely and surprising within an experiment. We did not find such enhancement effects in this situation, but instead observed repetition suppression, providing evidence against this hypothesis. Our results instead align with recent work reporting distinct mechanisms underlying repetition suppression and perceptual expectations (e.g. Grotheer & Kovacs, 2015; Pajani et al., 2017; Feuerriegel et al., 2018).

### 8.4. Caveats

A number of important caveats should be taken into account when interpreting our results. Firstly, although we manipulated face identity in our experiments, the repetition effects observed here largely index face *image* repetition rather than face *identity* repetition per se. This does not detract from our main findings, but means that oddball responses in our experiments should not be interpreted as examples of *facial identity discrimination* responses, as obtaining such responses was not the goal of this study. Experiments designed to assess facial identity discrimination with FPVS typically control for repetition of low-level image features by randomly varying size at every presentation cycle (e.g., Liu-Shuang et al., 2014; 2016; Dzhelyova & Rossion, 2014a; 2014b; Xu et al., 2017). Moreover, in these experiments different and highly variable face identities are introduced at every oddball cycle, rather than the same oddball face identity, preventing repetition effects across oddball changes, increasing the individual face discrimination response, and minimizing again the contribution of low-level visual cues to this individual face discrimination response (Liu-Shuang et al., 2014; 2016; Dzhelyova & Rossion, 2014a; 2014b; Xu et al., 2017).

Secondly, the results reported here were obtained in the context of an orthogonal task, a factor that conceivably may have inhibited detection of expectation/anticipation effects for face oddballs. Task-relevant fixation cross colour changes were unrelated to the face stimuli in our experiment, and expectation effects can be reduced or absent when attention is diverted to a different stimulus (Larsson & Smith, 2012; Hsu et al., 2014). We believe that expectation effects might indeed be detectable for repeated stimuli when faces are task-relevant (for example in a face identity detection task). However, this point is distinct from our finding that, when the critical stimuli are not task-relevant (as in most studies of VMRs), expectation effects are suppressed or abolished by immediate stimulus repetition.

Thirdly, the expectation effects observed in Experiment 2 could also be defined as delayed repetition effects. This is because there were many repetitions of the expected Identity Change oddball face images within a sequence (with each presentation separated by multiple base rate faces) compared to the surprising Identity Change oddball images in the same sequences. We believe that our results are primarily due to stimulus expectations rather than delayed repetition per se. This is because previous studies not using oddball designs, which did not have a delayed repetition confound, have reported similar interactions between expectation and immediate stimulus repetition (e.g., Todorovic & de Lange, 2012; Kovacs et al., 2012; Larsson & Smith, 2012; Amado et al., 2016). More generally, delayed repetition and stimulus expectation rely on very similar experimental manipulations in oddball designs (whereby perceptual expectations are driven by frequency of stimulus presentation). Distinguishing between these may require manipulations that test hypotheses regarding the underlying mechanistic implementations at the neural level (see Vinken & Vogels, 2017; Bell et al., 2017).

Finally, it is an open question as to whether the stimulus expectation effects observed in our study might extend to stimuli other than faces, for example simple visual stimuli (e.g., oriented bars) that are commonly used in visual oddball designs (e.g., Czigler et al., 2002; Tales & Butler, 2006; Kimura et al., 2009). Although there is some suggestion that stimulus expectation effects are reduced or absent for some object categories such as nonface objects and unfamiliar orthographic symbols (Grotheer & Kovacs, 2014; Kaliukhovich & Vogels, 2011, 2014; Kovacs et al., 2013), whether expectation effects do indeed exist for such non-face stimulus classes is beyond the scope of our study.

### 8.4. Conclusions

The research reported here indicates that visual mismatch responses are reduced with stimulus repetition. This finding uncovers a relationship between repetition suppression and perceptual expectation effects that has received little consideration in visual mismatch research thus far. Further investigation of this relationship will be critical to understanding the necessary conditions for visual oddball effects, and may reveal a hierarchy of interacting effects of expectation-based and stimulus exposure-dependent processes in the brain.

## Acknowledgements

D.F. was supported by an Australian Government Research Training Program Scholarship. This work was supported by an “Action de Recherche Concertee” grant (ARC; 13/18-053), the Louvain Foundation, and by a co-funded initiative by the University of Louvain and the Marie Curie Actions of the European Commission awarded to G.L.Q.

